# Dynamic modeling of EEG responses to natural speech reveals earlier processing of predictable words

**DOI:** 10.1101/2024.08.26.609779

**Authors:** Jin Dou, Andrew J. Anderson, Aaron S. White, Samuel V. Norman-Haignere, Edmund C. Lalor

## Abstract

In recent years, it has become clear that EEG indexes the comprehension of natural, narrative speech. One particularly compelling demonstration of this fact can be seen by regressing EEG responses to speech against measures of how individual words in that speech linguistically relate to their preceding context. This approach produces a so-called temporal response function that displays a centro-parietal negativity reminiscent of the classic N400 component of the event-related potential. One shortcoming of previous implementations of this approach is that they have typically assumed a linear, time-invariant relationship between the linguistic speech features and the EEG responses. In other words, the analysis typically assumes that the response has the same shape and timing for every word – and only varies (linearly) in terms of its amplitude. In the present work, we relax this assumption under the hypothesis that responses to individual words may be processed more rapidly when they are predictable. Specifically, we introduce a framework wherein the standard linear temporal response function can be modulated in terms of its amplitude, latency, and temporal scale based on the expectedness of the current and prior words. We use the proposed approach to model EEG recorded from a set of participants who listened to an audiobook narrated by a single talker, and a separate set of participants who attended to one of two concurrently presented audiobooks. We show that expected words are processed faster – evoking lower amplitude N400-like responses with earlier peaks – and that this effect is driven both by the word’s own predictability and the predictability of the immediately preceding word. Additional analysis suggests that this finding is not simply explained based on how quickly words can be disambiguated from their phonetic neighbors. As such, our study demonstrates that the timing and amplitude of brain responses to words in natural speech depend on their predictability. By accounting for these effects, our framework also improves the accuracy with which neural responses to natural speech can be modeled.

## Introduction

The human brain understands natural speech at rates of around 120–200 words per minute. A well-known signature of this process is the N400 electrophysiological brain response – a prominent centroparietal voltage negativity (N) at around 400ms after the onset of unexpected words. The N400 is typically revealed by recording and contrasting electroencephalogram (EEG) responses to many sentences with unexpected vs expected endings and then averaging the set of subtracted final word response profiles (Kutas & Hillyard, 1980). The time course of the N400 is very reliable and has been fleshed out over thousands of studies (Kutas & Federmeier, 2011). However, because the N400 is computed as an average of word response profiles, it may “blur” putative between-word differences in the timing of electrophysiological responses. There are several reasons to suppose these temporal differences could exist. From the perspective of efficient information processing, one might expect unsurprising words that convey little new content to be processed more rapidly to conserve neural energy and expedite comprehension. Indeed, years of research on the role of prediction in language (Ryskin & Nieuwland, 2023) suggests that unsurprising words are more easily identified (Fischler & Bloom, 1979) and read more quickly (Ehrlich & Rayner, 1981). Additionally, one might expect that the identification of a word might rely on the interaction between the word’s predictability and the word’s phonetic structure (more specifically, how quickly the word can be disambiguated from its phonetic neighbors). For example, although in general “at” is more frequent than “atmosphere”, the likelihood of encountering “at” is relatively reduced in the case of “CO2 in the Earth’s___“). Discovering such putative word/context-variant neural processing dynamics would be valuable for characterizing language processing in the brain, and predicting their occurrence could be key to advancing recent attempts to build accurate neuro-computational models of natural speech comprehension.

The notion of building computational models of natural speech and language processing in the brain is beginning to seem more tractable in light of recent advances in large-scale language modeling. EEG studies of natural speech comprehension have placed heavy focus on predicting the N400 using estimates of words’ unexpectedness derived from language models (Brodbeck et al., 2022; Broderick et al., 2018; Caucheteux et al., 2021; Heilbron et al., 2022). In particular language models, such as GPT-2, that are explicitly trained to predict next-word identity based on prior words, have led to natural speech N400 models based on so-called lexical surprisal estimates that quantify a word’s unexpectedness as the logarithmic inverse probability of that word coming next (N. J. Smith & Levy, 2013). These approaches have predicted EEG data by fitting linear regression mappings (known as temporal response functions or TRFs) (Crosse et al., 2016; Lalor et al., 2006) based on a series of lexical surprisal estimates that are time-aligned to word onsets and displaced at intervals to account for neural response delays (e.g. a displacement of +400ms would capture peak N400 response). The resultant profile of regression weights reliably traces the standard N400 response profile in centroparietal electrodes (Heilbron et al., 2022; Synigal et al., 2023). However, just like traditional contrastive EEG experiments, these linear mappings ultimately fit an average response profile across words and have no flexibility to capture temporal differences in neural responses to different words and contexts. The current study seeks to discover evidence for word/context-specific differences in neural response times by constructing new dynamically warped TRF models of EEG recordings of natural speech comprehension.

We hypothesized that the timing of the natural speech N400 electrophysiological responses will vary across stimulus words according to each word’s prior lexical context. To test this hypothesis, we implemented a new “dynamically warped TRF” model aimed at generating word/context-specific TRFs based on natural speech EEG data and then evaluated the model’s ability to predict new EEG data. The new dynamically warped TRF model estimates linear transformations of a canonical N400 TRF template – modulating TRF response latency, timescale and amplitude based on the lexical surprisal of each individual word in the stimulus. Lexical surprisal is computed using a modern, transformer-based large language model (GPT-2; Radford et al., 2019). We examined the new dynamically warped TRF model with EEG recorded from subjects who listened to an audiobook narrated by a single talker, and also subjects who attended to one of two concurrently and dichotically presented audiobooks. The results show that modulating TRF latency improves EEG prediction accuracy in the centroparietal channels that are traditionally associated with semantic processing of speech (Broderick et al., 2018; Heilbron et al., 2022; Kutas & Federmeier, 2011) and that this effect is specific to listeners paying attention to the modeled speech. In particular, we show that predictable words are processed expeditiously-eliciting early and low amplitude responses – and that the timing of a current word’s response is shaped by the lexical surprisal of both the previous and current words. Additional analysis demonstrate that this effect is not explainable simply on the basis of how quickly a word could be disambiguated from its phonetic neighbors.

## Results

### EEG recordings

We used two publicly available EEG datasets to test our hypotheses (Broderick et al., 2018). The single talker dataset was obtained from 19 participants listening to about 60 minutes of a narrative audiobook (split into 20 runs of about 3 minutes each). The cocktail party dataset was obtained from 33 participants listening to 30 minutes of two narrative audiobooks presented simultaneously – one in each ear – and with 16 participants attending to one story/ear and 17 participants attending to the other story/ear. Participants were presented with both audiobooks in 30 runs each of one-minute duration and asked to answer multiple choice questions on the content of both stories (Power et al., 2012). Each channel of the EEG recordings was referenced to the average data from two mastoid electrodes placed behind the ears, filtered between 0.5 and 8 Hz, interpolated if it was too noisy compared with the surrounding channels (see data preprocessing section below), downsampled to 64 Hz, and z-scored.

### Analysis overview

To test for differences in EEG response timing to different words in different (natural speech) contexts, we introduced computational methods to fit dynamically warped temporal response functions (TRFs) to individual words. Thus, rather than modeling neural responses to all words as simply scaling linearly with their lexical surprisal while having precisely the same temporal response profile, the new dynamic TRF has the flexibility to modulate the latency and time-scaling of TRFs for specific words in specific contexts, as well as fitting an additional scaling factor for their amplitude.

Specifically, the standard TRF assumes a linear relationship between changes in particular stimulus features and the resulting EEG response. This can be written as:

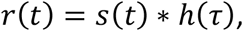

Where *s*(*t*) and *r*(*t*) indicate the stimulus feature and model-predicted neural response at time *t*, respectively, and *h*(τ) indicates the filter weight at time lag τ, relative to time *t* (please see Methods and Crosse et al., 2016 for more details). In our study – as we explain below – our primary stimulus feature of interest was the lexical surprisal of each word in context, represented as a series of impulses at word onset, with the impulse amplitude reflecting the surprisal magnitude.

While this approach has proven useful in a wealth of studies on the neural processing of speech stimuli (Ahmed et al., 2023; Brodbeck et al., 2022; Broderick et al., 2022; Heilbron et al., 2022), it explicitly assumes that the TRF, *h*(τ), has the same timing and same shape for every change in the stimulus feature. Of course, the approach allows for the response to change in amplitude – but it does not account for any effects that previous words might have on the response to the current word, and, again, it does not allow for any variations in the timing or shape of the response.

### Dynamically Warped TRF overview

To allow for more flexibility in the response, our approach here is to allow the TRF model to vary for every individual word. Specifically, we allowed the TRF for each word to vary in its timing and its shape – as well as its amplitude to scale in a word-specific way. To do this, we altered the TRF model equation as follows:

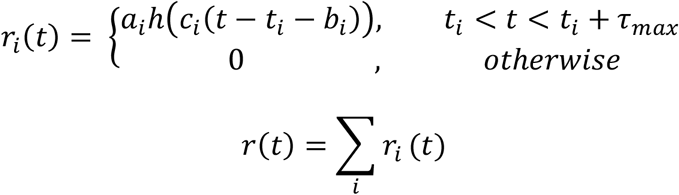

where *i* indexes each individual word, *t*_*i*_ indicates the onset time of the *i*th word, τ_*max*_ indicates the maximum value of time lag, *a*_*i*_ enables variations in the amplitude of the response of the *i*th word, *b*_*i*_ enables variations in the time shift of the response, and *c*_*i*_ allows for variation in the temporal scale (time-stretch) of the response. Importantly, the stimulus feature of interest, *s*_*i*_ – which in the present study was the lexical surprisal of the word at time *t*– is used to compute the transformation parameter (*a*_*i*_,*b*_*i*_,*c*_*i*_ ; as depicted in Figure 1). Here, we distinguish *s*_*i*_ with *s*(*t*) to highlight the fact that lexical surprisal was calculated for each word but not for each time point. It should be noted that we cropped both the original and time shifted TRFs to let them have the same start and end time.

**Fig. 1.**
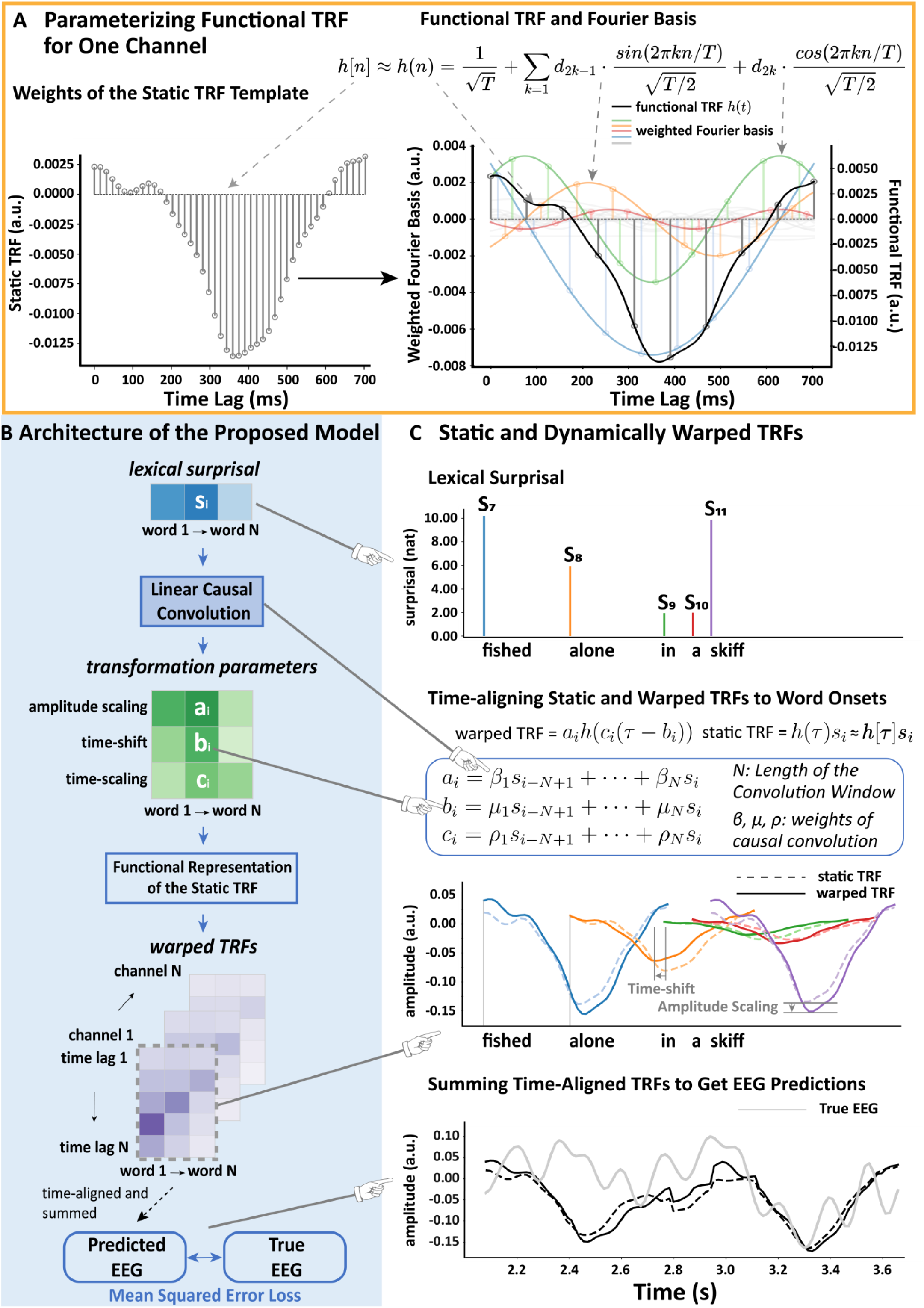
Overview of the approach. **A**, Representing TRF weights as a linear combination of Fourier basis functions. The left plot shows the discrete static TRF weights (canonical N400 TRF template) which were estimated using ridge regression. The right plot illustrates the functional representation of this discrete TRF template using continuous Fourier basis functions. The functional TRF that was parameterized by the static TRF is colored in black; the weighted Fourier basis functions are in other colors.**B**, The architecture of the proposed dynamically warped TRF model. The lexical surprisal values of each word in our stimuli were convolved with causal kernels to estimate three transformation parameters for each word (the number of kernels defines how many transformations the model can support). The obtained transformation parameters were then used to modify the functional TRF module to get a TRF specifically tailored to each word. Specifically, the TRF for each word, *i*, could be dynamically warped in terms of its amplitude, a_i_, latency (time shifting), b_i_, or temporal scale (i.e., time stretching), c_i_. Once these modifications had been made, the dynamically warped TRF for each word was aligned to that word’s onset and then summed across all words to enable prediction of the EEG response. The loss we used for training the causal convolutional layer was the mean squared error between the predicted EEG and the recorded EEG.**C**, Illustration and comparison of static and dynamically warped TRFs estimated for words picked from the single-talker stimuli. It shows how the lexical surprisal values (top), and the TRFs (middle) were aligned to word onsets. The predicted EEG (bottom) was obtained by pointwise summing the time-aligned TRFs across words. The equation in the middle specifies how lexical surprisal was convolved with the causal kernels to get transformation parameters shown in B (green grids) for transforming the canonical N400 TRF template shown in A (left).

By allowing the TRFs to be modulated in this way, our overarching goals were: 1) To test if dynamic TRFs provide a more accurate EEG model than the standard static TRF approach. This would provide evidence for context and word-specific neural response timing. 2) To explore the relationship between EEG response timing and lexical surprisal, with the hypothesis that unsurprising words are processed more rapidly.

The dynamically warped TRF (Figure 1B, 1C) implements word/context-specific linear transformations of the standard static TRF. As such, the dynamic TRF begins with the computation of a static TRF – as a predictive multiple regression mapping from time-lagged stimulus features to individual electrode responses. Given that we were interested specifically in how the unexpectedness of individual words affects EEG responses, we focused our analysis on TRFs derived from the lexical surprisal of each word. Specifically, we fit static TRFs to EEG using a time series of GPT-2-based lexical surprisal impulses placed at word onsets. As we have done in previous work (Broderick et al., 2022), we also included several other predictors in an effort to explain additional variance in the EEG that might derive from features that are correlated with the lexical surprisal predictor. This included one feature controlling for lexical processing, and another controlling for lower-level speech acoustics. The lexical control feature was simply a train of unit impulses marking the onset of each word. The low-level speech acoustic feature was the audio envelope of the speech stimuli. All three features were time-lagged between 0 and 700 ms. Static TRFs were fit using ridge regression in a nested cross-validation framework (see Cross Validation section). Consistent with previous research (Broderick et al., 2018; Synigal et al., 2023), this static TRF resembled the classic N400 event-related potential – with a marked negativity at a timelag of ∼400 ms (Fig 1A).

The idea then was to warp this static TRF (in amplitude, latency (time-shift), and temporal scale (time-scaling)). To do this, we needed to learn the amplitude, time-shift and time-scaling parameters for every word. We chose to do this by incorporating a linear convolutional layer into our framework – where we could use the error between EEG predictions and recorded EEG to learn the three parameters of interest. This involved backpropagating this error through the network to update the weights of that network (i.e., the weights that generate our three parameters of interest). To make this possible, we first represented the static TRF in terms of continuous functions (e.g., a Fourier basis set; Fig 1A) such that it would be differentiable (allowing us to estimate error gradients and minimize our loss). Specifically, we wanted to represent the TRF on each EEG channel as the linear sum of several Fourier basis functions that would capture the overall shape of the original static TRF (Fig 1A). We did this for each individual channel separately – prior to network training – via Functional Data Analysis (Ramsay & Silverman, 2005). The Fourier basis weights for each of the 128 EEG channels were then held constant for all subsequent analyses (including network training), and these basis functions could then be scaled by the three learned parameters. These three parameters were fed into the functionally represented TRF to modulate the TRF amplitude, latency (relative to word onset), and temporal scale. To reduce model complexity precisely the same three parameters were used for each of the 128 channels – thus our current model assumes that TRFs vary across different words but that this variation is shared across electrode sites.

The TRF amplitude, time-shifting, and time-scaling parameters were derived from the weighted outputs of a linear causal convolution layer – which accordingly consisted of three nodes (one for each parameter). Each node performed a linear causal convolution over an ordered sequence of lexical surprisal values, with one value per word (not per timepoint). In practice, we found that convolution over only the current and preceding lexical surprisal values was sufficient to achieve maximal EEG prediction accuracy (see later analyses in Supplementary Fig 1). Consequently, the dynamic TRF’s input layer contained two nodes representing the two consecutive surprisal values (unless stated otherwise) which fed into the hidden convolutional layer. Importantly, given the challenge of fitting three parameters for each word, we constrained our analysis to derive the same three parameters for all EEG channels and for all participants. In other words, we fit our parameters in a single procedure that included all channels and participants.

To predict new EEG data, we constructed a time series populated by predicted TRFs by: (1) Estimating time-shifted and time/amplitude-scaled TRFs for each word. (2) Time-aligning each warped TRF to the corresponding word onset in a new time-series that otherwise contained zeroes. (3) Pointwise summing all new time-series across words from (2) to produce a single predicted time-series. Steps (1-3) were repeated for each of the 128 channels. To combine the new dynamically warped lexical surprisal time-series with word onset and speech envelope control predictions: (1) Time-series reflecting EEG responses to control features were predicted via a matrix multiplication of the word-onset spikes and speech envelope with the corresponding weights of the pre-computed static TRF mapping (which also supplied the N400 template). (2) The predicted control time-series were pointwise summed with the new dynamically warped lexical surprisal time-series to generate a single predicted time-series. This was repeated for each electrode. [For more details of the process used to optimize the weights of the causal convolution layer – and thus to learn the transformation parameters – please see the Methods.]

### EEG responses to expected words peak earlier, but do not last longer than for surprising words

Using the single talker dataset, we first tested the hypothesis that modeling EEG with dynamically warped lexical surprisal TRFs would lead to more accurate predictions of new EEG data (without considering phonetic information for now). We reasoned that establishing improvements in EEG prediction accuracy would provide evidence that the timing of brain responses varies according to lexical context. Then, inspecting the dynamic model weights can help expose the relationship between lexical surprisal and the timing of brain responses. To recap, we had further hypothesized that unsurprising words that add little new content would be rapidly processed.

To estimate the predictive contribution of dynamically time shifting the TRF and/or scaling the TRF timescale and/or amplitude, we ran a battery of comparative analyses that modeled and predicted audiobook EEG data with the dynamically warped TRF, as well as reduced variants that were trained with one or two transformation parameters missing (e.g., with scale factor or amplitude scaling removed). A first observation was that we saw no predictive benefit to dynamically time-scaling the TRF (W = 19, p = 0.9995, single-tailed). As such, we simplified the following presentation of results to focus only on TRF time-shift (Time), amplitude scaling (Amp) and their combination (Time&Amp).

To evaluate whether the dynamically warped TRF approach yielded more accurate EEG predictions than the static TRF, we deployed signed ranks tests to compare respective EEG prediction accuracies both on individual electrodes (Fig. 2 A), and across the entire scalp, i.e., by averaging prediction accuracies across all 128 electrodes. All EEG predictions reflected the combination of lexical surprisal with the word onset and speech envelope control features. Assessing prediction accuracy on individual channels revealed that the dynamic TRF outperformed the static TRF on a substantial number of individual electrodes (Fig 2A, Right), and that those electrodes were similar to the electrodes that show good predictive power for the static surprisal model (Figure 2A, Left). Averaging across the entire scalp (Fig 2B), we found that dynamically time shifting the TRF (Time) yielded more accurate EEG predictions than the static TRF approach (Scalp-average signed ranks: W = 147.0, p = 0.0180, single-tail). Incorporating scaling of the TRF amplitude along with the time shift (Time&Amp) yielded stronger predictions still (scalp-average W=169, p=0.0008) over the static TRF.

**Fig. 2.**
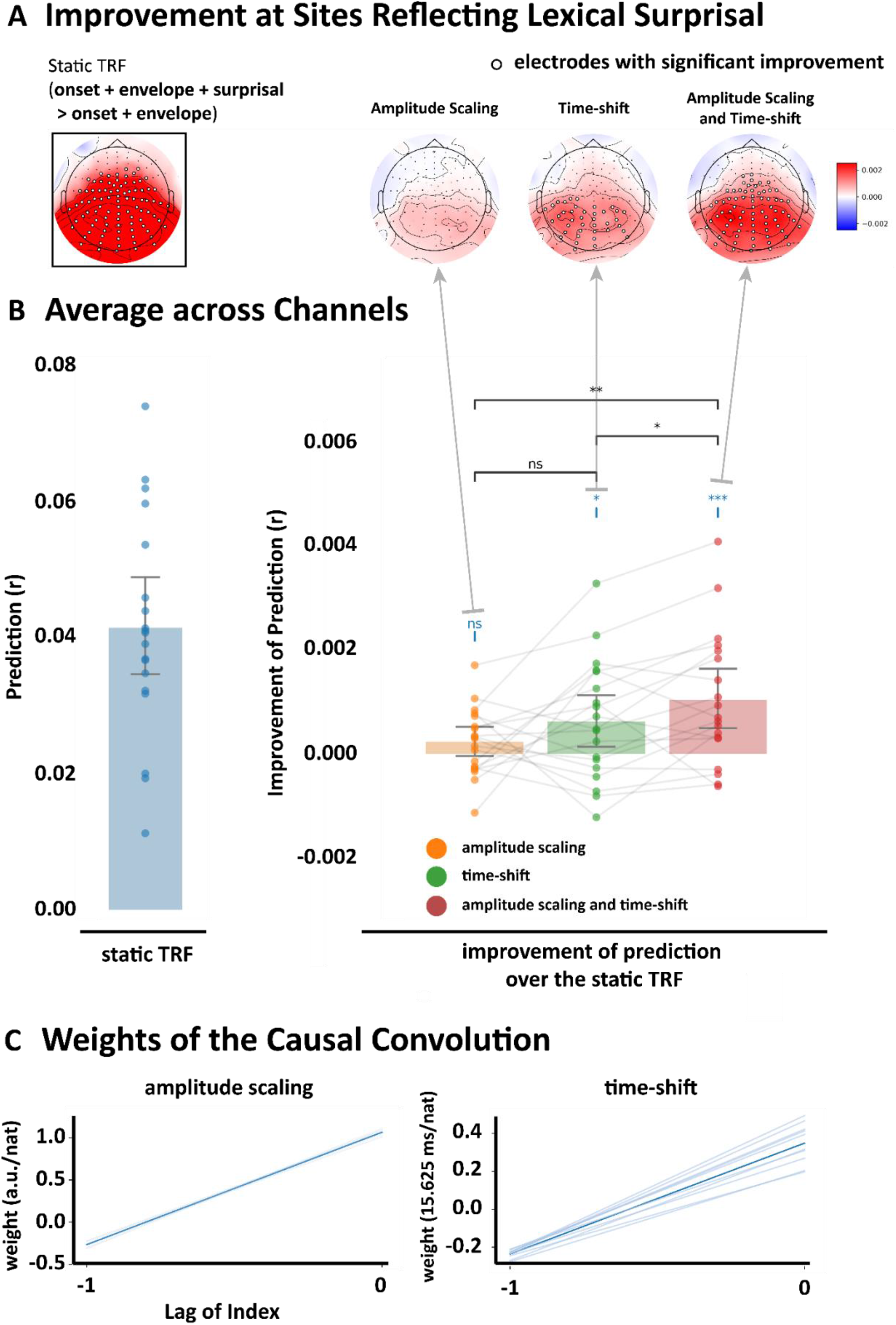
Time-shifted TRF improves speech prediction accuracy. The prediction accuracy was measured using Pearson’s correlation, and the significance of improvement was measured using paired Wilcoxon signed rank tests across participants. Electrodes where the prediction improvement is significant (p < 0.05) were labeled with white circles. P values were corrected using false discovery rate based on Benjamini-Hochberg correction (FDR), with alpha set to 0.05.**A**, Topographic maps for comparing prediction accuracies of different TRF variants on individual electrodes. The left map (in the black box) shows the improvement of prediction accuracy for a static TRF when adding lexical surprisal as a predictor in addition to the acoustic envelope and onset control predictors. The right three maps then show the additional improvement of three dynamically warped TRF variants over the static TRF. From left to right, the three variants are amplitude scaling (Amp), time shifting (Time), and their combination (Time&Amp).**B**, The bar plot quantifies prediction accuracies of the same static TRF (left) and dynamically warped TRFs (right) averaged across all 128 channels. Text/asterisks in blue in the right panel indicate statistical comparison results between each dynamic TRF and the static TRF. Horizontal connecting bars indicate the comparison between the different dynamic TRF variants. Significance is indicated by * if p < 0.05, ** if p < 0.01, and *** if p < 0.001.**C**, Convolutional kernel for the amplitude scaling (left) and time-shift (right) parameters. The lag indicates the index relative to the current time step (e.g., 0 and -1 indicate the weights for the current and previous word, respectively). Nat indicates the natural unit of information of the surprisal values. Light blue colored lines indicate the weights of models fitted from individual rounds of cross-validation (each round of which involved training an all but two runs from every participant – i.e., for each participant, two runs were left out for testing). The deep blue colored line indicates the weights averaged across those cross-validation rounds.

To then interpret the relationship between response timing and lexical surprisal (in light of the hypothesis that the brain processes expected words more rapidly), we examined the convolutional weights linking lexical surprisal input values to the time-shift node in the Time&Amp model (see Fig. 2C). Time-shift weights for the current lexical surprisal value were positive, which indeed suggests that electrophysiological responses to less predictable words (higher surprisal) peaked later than those to expected words. In contrast, time-shift weights on the previous lexical surprisal value were negative with lower magnitude. This suggests brain responses to the current word may peak earlier if the *previous* word was surprising. A similar pattern of positive/negative weightings across current and prior lexical surprisal values was observed for amplitude scaling. Thus, if there were two surprising words in a row, the response to the latter may be earlier and shallower than if the preceding word had instead been unsurprising.

Interestingly, this interaction between the lexical surprisal of the current and previous words was crucial to the improved prediction accuracy of the dynamic TRF in the single talker condition. In particular, a dynamically warped TRF based on the lexical surprisal value of the current word alone produced no improvement in prediction accuracy over the static TRF (W = 124, p = 0.1289; Supplementary Fig. 1). And, incorporating the lexical surprisal of more preceding words did not lead to any systematic additional improvement.

In sum, this section provides evidence that (1) dynamically warped TRFs can provide more accurate EEG models than static TRFs, (2) the human brain expedites processing of expected words, and (3) that the preceding word’s likelihood influences the timing and amplitude of brain responses to the current word.

### Dynamically warping TRFs provide more accurate models of EEG responses to attended but not unattended speech

In the previous section, we provided evidence that dynamically warping the amplitude and time shift of a TRF based on lexical surprisal can provide more accurate EEG predictions than the original static TRF. To confirm that this result was driven by differences in brain responses when comprehending expected/unexpected words – and not by potentially spurious correlations with the acoustics of expected/unexpected words (Aylett and Turk 2004; Bell et al. 2009; Lieberman 1963) – we applied the same procedure as above to a cocktail party attention dataset. The cocktail party attention dataset was collected during an experiment wherein participants were asked to attend to one of two concurrently and dichotically presented audiobooks (O’Sullivan et al., 2015; Power et al., 2012). We hypothesized that dynamic TRF warping would produce no improvement in performance for EEG responses to unattended speech. This hypothesis was based on previous work that found little to no evidence of contextual semantic processing for unattended speech (Broderick et al., 2018).

We applied the same dynamic TRF procedure – and comparison with the static TRF approach – to EEG data modeled based on both attended and unattended speech. We again chose to focus our analysis on the effect of modifying the TRF in Time, in Amp and in both Time&Amp, because we continued to observe no predictive benefit as a result of dynamically scaling the TRF timescale (attended: W = 87, p=0.9998, single-tail; unattended: W = 229, p=0.821, single-tail).

We first found that the Time&Amp dynamic TRF in the attended condition yielded stronger predictions than the static TRF at a substantial number of central electrodes (Fig 3A, Left), consistent with it being an improvement in modeling responses based on lexical surprisal (since the distribution is similar to the classical N400-like topography). The dynamic TRF did not outperform the static TRF at any individual electrode for unattended speech (Fig 3A, Right). These findings were also reflected in prediction accuracies averaged over the entire scalp, with a significant improvement for the Time&Amp dynamic TRF for attended speech (W = 401, p=0.0154, single-tail; Fig 3B), but not for unattended speech (W = 176, p=0.9699, single-tail). In addition, directly comparing the benefit of using the Time&Amp dynamic TRF over the static TRF for attended speech with that for unattended speech revealed that the benefit was significantly greater for a substantial number of channels (Fig 3C).

**Fig. 3.**
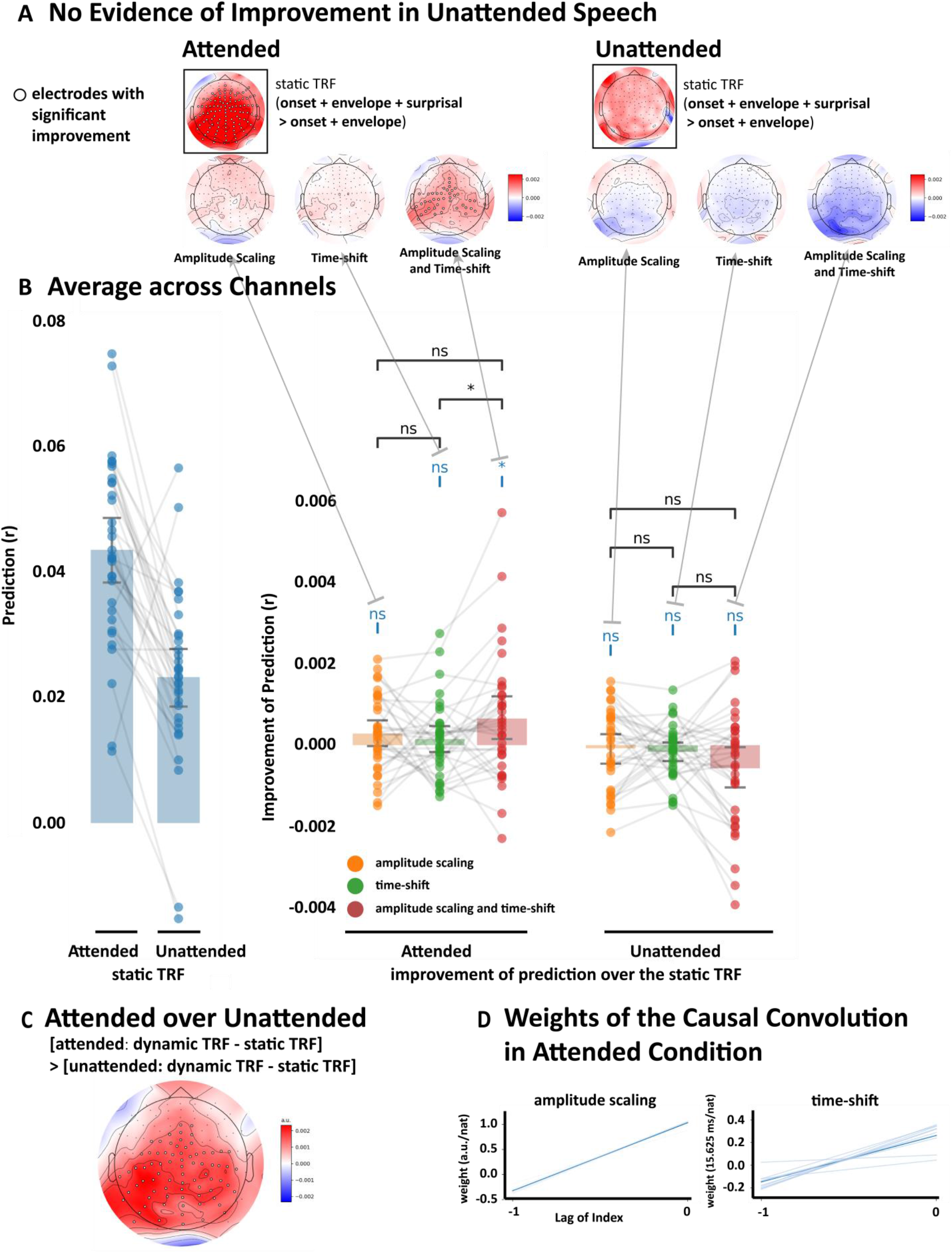
The dynamic TRF improves prediction accuracy significantly only for attended speech. The prediction accuracy was measured using Pearson’s correlation, and the significance of improvement is measured using paired Wilcoxon signed rank tests. Electrodes where the prediction improvement is significant (p < 0.05) were labeled with white circles. P values were corrected using false discovery rate based on Benjamini/Hochberg correction (FDR), with alpha set to 0.05.**A**, Topographic maps for comparing EEG prediction accuracies using different variants of the dynamic TRF. Same as Figure 2A, maps in the black box quantify the electrodes where including lexical surprisal as a predictor leads to significant improvements in EEG prediction under the static TRF in attended and unattended speech, respectively. Other maps on the top quantify the sites where different variants of dynamic TRF have improvement over the static TRF in both conditions.**B**, Bar plots for comparing EEG prediction accuracies using different variants of the dynamic TRF. Same as Figure 2B, the blue bar plots (left) show the prediction accuracy of the fixed TRF for attended and unattended speech. The other bar plots (right) show the improvement of prediction accuracy across the entire scalp (i.e., averaged across all channels).**C**, Directly comparing the benefit of using the Time&Amp dynamic TRF over the static TRF for attended speech with that for unattended.**D**, The weights of the convolutional layer for the amplitude scaling and time shifting. The lag of index indicates the index relative to the current time step of convolutional weight (e.g., -1 indicates the weight for the last word and 0 indicates the weight for the current word). Nat indicates the natural unit of information. Light blue colored lines indicate weights of models from individual folds of cross-validation. The deep blue colored line indicates the averaged weights across all the models.

Similar to single talker results, weights from the attended condition were positive and larger for the current lexical surprisal value, and smaller and more negative for the previous lexical surprisal value – both for amplitude scaling and time shifting (Fig 3D). This again indicates that more surprising current words lead to later and larger responses, with more surprising previous words reducing those effects. As was the case for the single talker dataset, the interaction between the lexical surprisal of the current and previous words was essential for the improved predictive power of the dynamic TRF (Supplementary Fig. 1). Fitting such a dynamically warped TRF based on the lexical surprisal value of the current word alone produced no improvement in prediction accuracy over the static TRF for the attended speech (W = 242, p = 0.7545) or (perhaps trivially) the unattended speech (W = 263, p = 0.6245). One caveat here: we saw improved predictions for the single talker condition when using dynamically warped TRFs based on the current and previous word (i.e., with a window length of 2), but also for window lengths of 3, 4, 7 and 9. For the attended speech condition, we only saw the effect for a window length of 2. As such, the result must be interpreted with some caution. The fact that the pattern of weights was so similar for the attend speaker condition and the single talker condition gives us some confidence in the finding. It is also worth noting that we found no significant effect for any window length for the unattended speech. It may be that the signal-to-noise ratio of the TRFs in the cocktail party data was lower than that for the single speaker data, meaning we only saw the effect for a window length of 2.

To sum up, this section corroborates the results of the last section, by showing that the dynamically warped TRF provides a more accurate model of EEG responses to natural (attended) speech. Secondly, it reveals that this improvement is selective to attended speech, and is not detectable when unattended speech is modeled and the brain disengages lexical processing.

### Time shifting and amplitude scaling covary with lexical surprisal

In the last two sections, we observed similar profiles in kernel weights assigned to time shifting and amplitude scaling (Fig. 2C, Fig. 3B) for (attended) speech. In both cases, the current surprisal value has a strong positive weight and the previous surprisal value has a weaker negative weight. This suggested that the variation in both time shifting and amplitude might be similar, and therefore that our network might be simplified by reducing the convolutional layer to a single variable modulating both time shifting and amplitude scaling. Because time shifting and amplitude modulation operate on different scales, the model consisted of a single convolutional operation, whose amplitude was then separately scaled for each output using a learned scalar variable (optimized in the same way as the convolutional kernel).

We hypothesized that the simplified model would be no less accurate in predicting single talker and cocktail party EEG data. This would support the idea that the time shifting and amplitude scaling effects are correlated and, thus, carrying similar information. Indeed, we found that there was no significant difference in EEG prediction accuracy between the original Time&Amp model and the shared-weights variant for either the single talker (N = 19, W = 94, p = 0.5235) or the attended speech (N = 33, W = 317, p = 0.2625) data.

To confirm the shared convolutional layer also has a similar pattern of weights to the results shown in previous sections, we examined the weights of the shared causal convolutional layer (Fig. 4A). Similar to the pattern in the previous sections, the weights estimated for both the single talker and attended speech conditions had a negative but smaller weight for the prior lexical surprisal value, and a larger and positive weight for the current lexical surprisal value. We also examined the relationship between surprisal and time shift for each word directly (Fig. 4B). The time-shift values were obtained by applying the fitted Time&Amp models on the testing data within each round of cross-validation. We also visualized how the time shift and amplitude of the dynamic TRFs varied with surprisal (Fig. 4C) by randomly selecting some words along the regression line (labelled points in Fig. 4B). For the single talker dataset, we visualized responses on midline parietal channel Pz, and for the cocktail party dataset, we visualized responses on the midline central channel Cz channel. Based on visual inspection of Fig. 4 (B, C), it is clear that both the time shiftand amplitude of the responses are larger when the lexical surprisal value of the current word are larger.

**Fig. 4.**
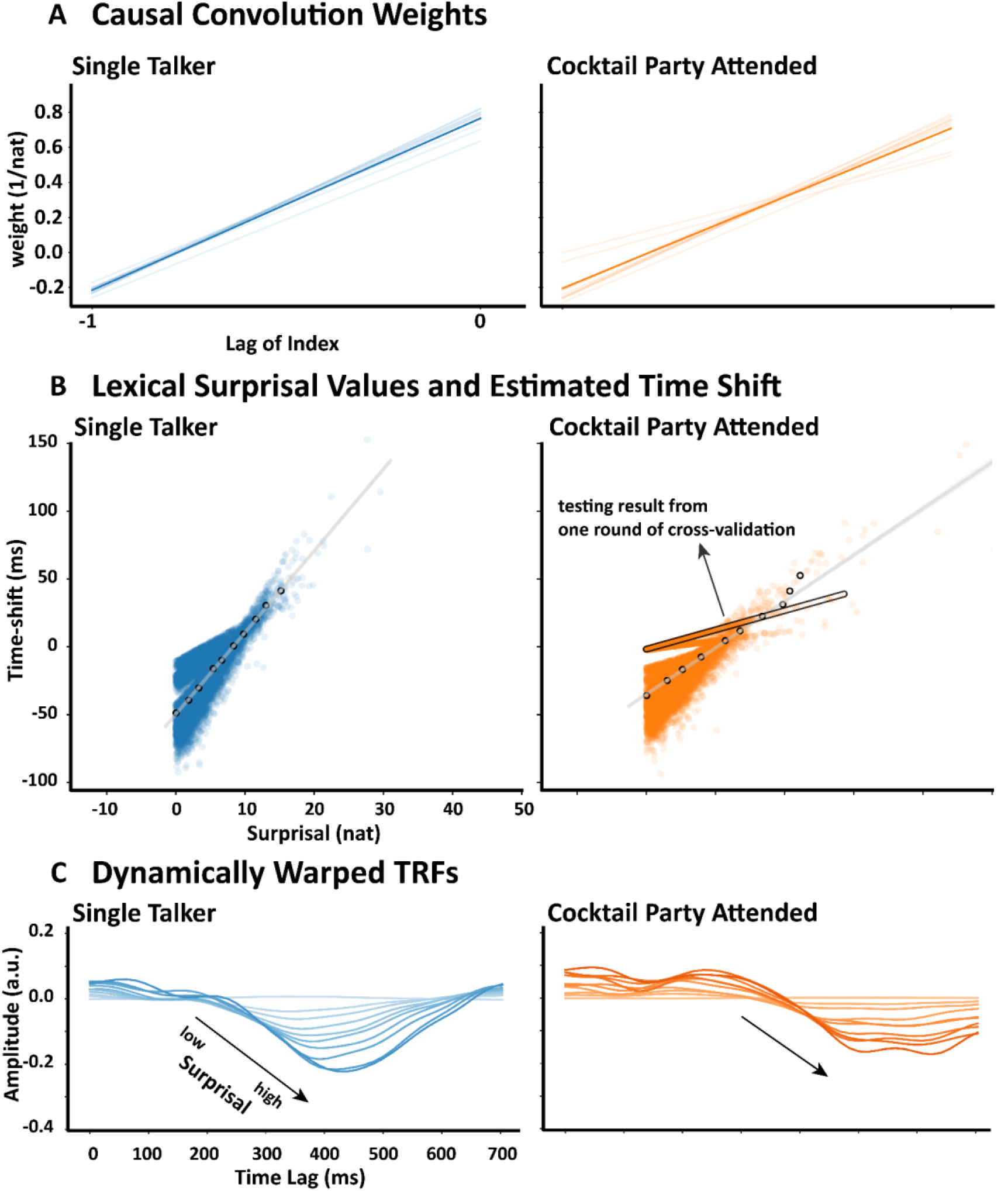
Time shifting and amplitude scaling covary with lexical surprisal.**A**, The weights of the convolutional layer shared by both time shifting and amplitude scaling. The lag of index indicates the index relative to the current time step of convolutional weight (e.g., -1 indicates the weight for the last word and 0 indicates the weight for the current word). Nat indicates the natural unit of information.**B**, The scatter plot of the transformation parameters. Each point indicates a pair of estimated time-shifting and amplitude scaling parameters for a word from all the testing data within each round of cross-validation. A clear positive relationship is visible for the single talker and attended cocktail party speaker.**C**, Visualization of time-shifted and amplitude scaled dynamic TRFs corresponding to the selection of words indicated in **B** (the white dots). The intensity of color varies according to the value of time shift (which is equivalent to using the amplitude since they share the same output node of the convolution layer).

### The relationship between lexical surprisal and EEG response latency is not explained by phonetic distinguishability

As mentioned in the introduction, we were also interested in the possibility that the identification of a word might be related to how quickly that word could be distinguished from its phonetic neighbors and, indeed, how this property might influence brain responses to a word beyond any effects of the word’s predictability. To explore this, we initially considered examining how a static TRF might be dynamically warped based on each word’s so-called uniqueness point (i.e., the phoneme in the word where that word diverges from all other words in the language; Marslen-Wilson, 1991). However, on calculating this uniqueness point for each word, we discovered that it was strongly correlated (0.504 on average across all stimuli used in this dataset) with lexical surprisal (this is because longer words tend to be more informative on average; Piantadosi et al., 2011). This made it difficult to disentangle the separate influences of predictability and uniqueness point on word identification. Indeed, it also caused us to worry that the lexical surprisal effect reported above might be more simply explained based on the uniqueness point of each word. To test this possibility, we built two additional models.

In the first model, we simply replaced lexical surprisal with the uniqueness point of each word as the input to each of the three nodes in the convolutional layer in the framework described above. (The uniqueness point was computed by identifying the time relative to offset of the first phoneme that enabled the word to be uniquely identified.) The idea here was that, if the uniqueness point of words explains the lexical surprisal effects we see on the dynamic TRF, then we should see very similar effects (including improved EEG prediction accuracy) when using uniqueness point as input. This turned out not to be the case. Specifically, we used signed rank tests to compare the EEG prediction accuracies between the uniqueness-point Time&Amp model and its corresponding static model, both on individual electrodes (Fig. 5B Uni) and across the scalp (by averaging predictions across all channels; Fig. 5A red bars). We found that the uniqueness-point Time&Amp model performed worse in both the single talker condition (scalp-average W = 9.0, p = 0.9999) and the attended speech in the cocktail party experiment (scalp-average W = 139, p = 0.9950). This result indicates that the uniqueness point is not the driving force behind the lexical surprisal effect reported in previous sections.

**Fig. 5.**
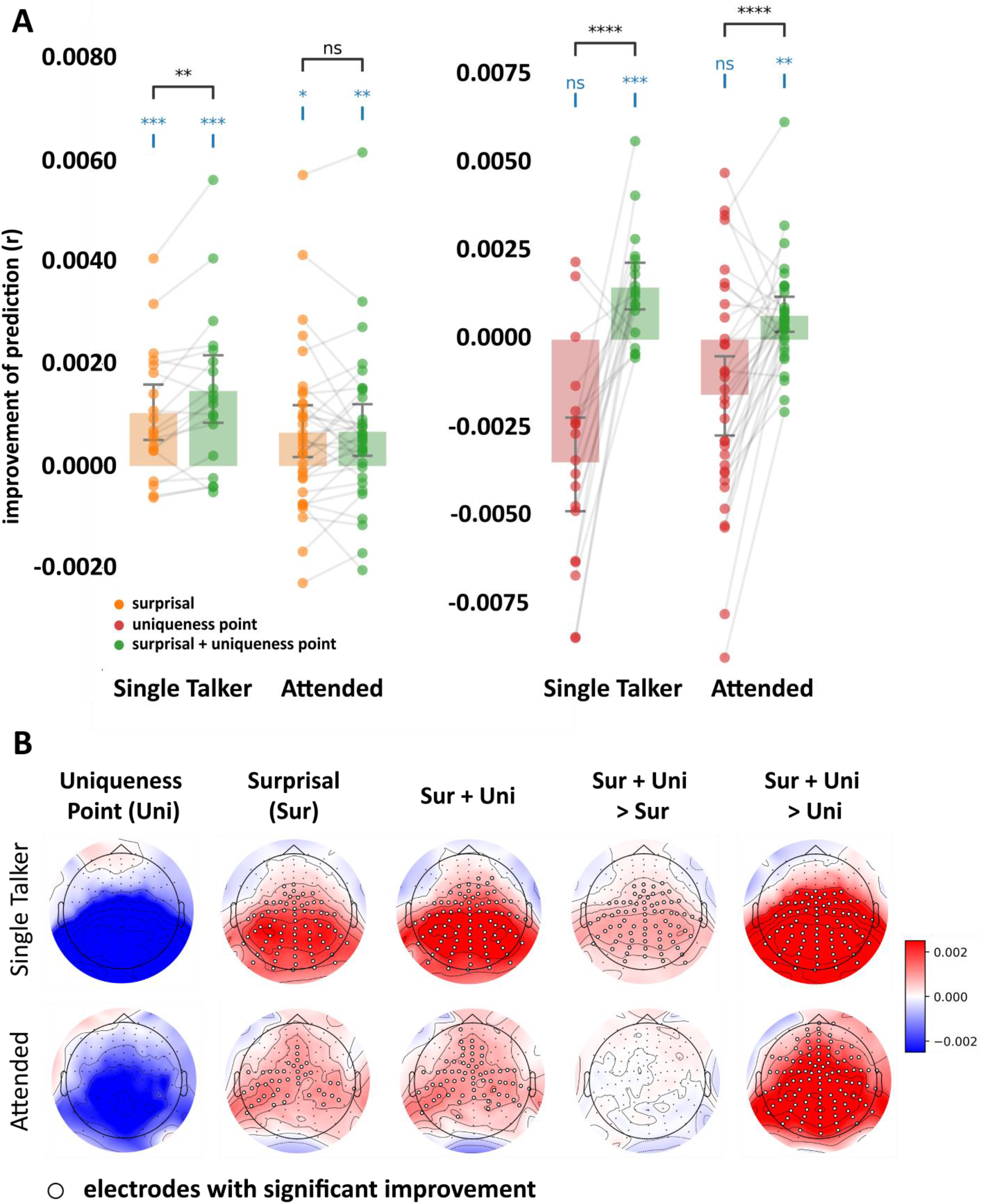
Lexical surprisal has a similar impact on N400 TRF latency and amplitude when controlling for the phonetic uniqueness point of words.**A**, Bar plots comparing the improvement in EEG prediction accuracy – relative to the static TRF – of the Time&Amp dynamic TRF models that are modulated by surprisal (orange), uniqueness point (red), or a linear combination of both (green). The uniqueness point indexes the timepoint at which a word becomes phonetically disambiguated from other competing words. Note that the green bars in the left and right bar plots are referenced to the same data.**B**, Topographic maps for the improvement of the Time&Amp dynamic TRF with surprisal (Sur), uniqueness point (Uni), or the linear combination of both (Sur + Uni) over the static TRF (the left three columns); and the differences between the Sur&Uni and the Sur dynamic TRF, or between the Sur&Uni and the Uni dynamic TRF (the right two columns).

In the second model, we added the uniqueness point as a second input channel – along with lexical surprisal – to each of the three nodes in the convolutional layer in the framework described above. Thus, in this augmented model, each node in the convolutional layer received two inputs, with each input convolved with a 2-tap convolutional kernel: the current and previous phonetic uniqueness point, and the current and previous lexical surprisal values. We again included word onset and acoustic envelope as co-predictors. And the model was then trained and deployed precisely as described for lexical surprisal above. The idea here was to enable us to test if the prediction accuracy of this augmented model (which included both lexical surprisal and the uniqueness point of words) was better than the prediction accuracy of the first model (which included only the uniqueness point of words). Such improved performance would be further evidence that lexical surprisal is driving the effects on the dynamic TRF, independent of the uniqueness point of words. This turned out to be the case. Specifically, we used signed rank tests to compare the EEG prediction accuracies between the augmented model and the uniqueness-point only model, both on individual electrodes (Fig. 5B Sur + Uni > Uni) and across the scalp (by averaging predictions across all channels; Fig. 5A right). We found that lexical surprisal improves the prediction performance significantly in both the single talker condition (scalp-average W = 181.0, p = 0.0000), and the attended speech (scalp-average W = 482.0, p = 0.000). This result shows that the lexical surprisal effects are not entirely explainable based on phonetic uniqueness point.

Finally, for completeness, we also sought to evaluate the effect of adding the phonetic uniqueness point information beyond the original lexical surprisal only model. Specifically, we used signed rank tests to compare the EEG prediction accuracies between the surprisal Time&Amp model and the augmented model that included uniqueness point, both on individual electrodes (Fig. 5B Sur + Uni > Sur) and across the scalp (by averaging predictions across all channels; Fig. 5A left). We found that the phonetic-augmented Time&Amp model performed better in the single talker condition (scalp-average W = 160.0, p = 0.0036). This supports the possibility that lexical surprisal and the uniqueness point of a word both influence the latency and amplitude of the brain response. However, when looking across the entire scalp, this result did not replicate for the attended speech in the cocktail party experiment (scalp-average W = 300.0, p = 0.3688), perhaps as a result of a slightly lower SNR in the EEG data from a multispeaker experiment. Based on visual inspection of Fig. 5B, it is clear that, in the single talker condition, the uniqueness point-augmented model yielded better EEG predictions on centro-parietal electrodes (Fig. 5B top right), which are the same electrodes that reflect lexical surprisal using the standard static TRF (Fig. 2A left). Although there were no such predictive benefits in the attended condition across the entire scalp, some individual electrodes over centroparietal scalp showed a significant improvement after adding the phonetic uniqueness point information (Fig. 5B middle right).

In sum, lexical surprisal adds significant value to the dynamic TRFs even when accounting for the uniqueness point of words. Considering the uniqueness point of words alone does not add predictive value to the dynamic TRF beyond the static (surprisal) TRF – although the partial improvements seen when adding both lexical surprisal and uniqueness point of words as inputs appears to provide some weak support for the idea that both features contribute to influence the EEG response to words. However, we do not wish to be too strong on the contribution from uniqueness point – given that it failed to add predictive value on its own and that it is significantly correlated with the lexical surprisal, which we know influences the responses.

## Discussion

Our study suggests that when listening to natural speech, the response to the lexical surprisal of a word, based on its preceding context, is not rigidly time-locked to word onset. By allowing a traditional static temporal response function to vary nonlinearly in terms of its time shift and its amplitude (Time&Amp), we found that both the amplitude and latency of the response to a word are influenced by the predictability of that word and the immediately preceding word. We demonstrated these finding by showing that a Time&Amp dynamic TRF model could better predict EEG collected under different conditions, relative to a standard, static TRF. The results show that, (1) dynamically warped TRFs can provide more accurate EEG models for lexical surprisal values than static TRFs, (2) the human brain expedites processing of expected words by reducing both the response latency and response amplitude, (3) the preceding word’s likelihood influences the timing and amplitude of brain responses to the current word in an opposite direction. We also confirmed that this result was not explainable simply as a result of how quickly a word could be distinguished from its phonetic neighbors.

As with any effort to optimize a model based on its ability to predict data, our dynamic TRF approach runs a risk of overfitting. That said, we used cross-validation as one approach to mitigate this concern. Furthermore, we developed the approach on a single-talker dataset and then validated that that approach performed similarly for only the attended speaker in a cocktail party dataset. Indeed, qualitatively, the analysis of both the single talker and the attended cocktail party talker produced similar results in terms of: 1) improved EEG modeling accuracy; 2) effects of lexical surprisal on response amplitude and latency; and 3) the fact that the responses to words are influenced in opposite directions by the surprisal of both the current and preceding word. The fact that this pattern of results did not hold for unattended speech is additional support for the idea that this consistent pattern of findings is not simply based on overfitting. That said, absence of evidence is not evidence of absence. It may be that the lower signal-to-noise ratio of responses to unattended speech has meant that the dynamic TRF simply could not be fit well to weaker responses in that condition. Indeed, we might go so far as to say that – based on previous research – responses to the context-based lexical processing of words in an unattended speech stream tend to be close to zero (Broderick et al., 2018).

Our analysis of the pattern in the convolutional weights for single talker and attended cocktail party speech revealed that those weights were positive for the lexical surprisal of the current word and negative with lower magnitude for the lexical surprisal of the previous word. This indicates that, when the current word is difficult to predict, it produces a larger, later response. However, this effect is reduced when the previous word was also difficult to predict. Indeed, a dynamically warped TRF based on the lexical surprisal value of the current word alone produced no improvement in prediction accuracy over the static TRF (Supplementary Fig. 1). This suggests that the influence of current word’s surprisal on the EEG depends on the surprisal of the previous word – with the time delay and amplitude increase being most pronounced when there is a relatively large surprisal for the current word after hearing a previous word that was not surprising. In order to learn this relationship, our model needed access to the surprisal values for both the current and previous word. This might explain why a relationship between word predictability and response latency has not previously been reported for the classic N400 ERP component – because the focus of those studies is typically on the predictability (cloze probability) of a single word. That said, the relationship we found between surprisal values and EEG response amplitude is consistent with the fact that the amplitude of the classic N400 tends to diminish with repeated presentations of surprising words (Petten et al., 1991).

A second initial goal of our study was to explore the possibility that, due to the phonetic structure of language itself, we might expect different words to be recognized at different latencies (e.g., consider “at” vs “atmosphere”). For example, it has been shown that listeners are faster in deciding whether a spoken item is a word (vs. a nonword syllable) when its uniqueness point is early in the word (Marslen-Wilson, 1991). This was not straightforward in our dataset because of a strong correlation between the uniqueness point and the lexical surprisal of each word. However, a series of analyses suggested that our primary finding of an influence of lexical surprisal on the timing and amplitude of EEG responses to words could not be explained as a result of correlations with the uniqueness points of words. Specifically, providing the uniqueness point of words as a lone input to the dynamic TRF pipeline did not improve EEG response predictions beyond those based on a static TRF. Moreover, dynamically warping TRFs based on using the combination of both lexical surprisal and the uniqueness point of words significantly outperformed a model based only on the uniqueness point of words. However, we also found that a combined model outperformed a model based only on the lexical surprisal of words for one of our data sets. This suggests the possibility that the two lexical properties might both influence EEG responses to speech. That said, the fact that a dynamic TRF based on the uniqueness point of words was no better than a static TRF means that we remain circumspect on this last point. Again, because these two lexical features were correlated, interpreting these nuanced results is not straightforward. Future experiments – likely involving speech stimuli that are constructed to decorrelate the lexical surprisal of words from their uniqueness point – will be required.

Although the proposed variable TRF model provides an easy-to-interpret tool for modeling the dynamic nature of responses to speech, there are a number of ways in which it can be improved in future work. First, given the need to derive a functional representation of the TRF (so that the neural network can perform gradient descent), the analysis is based on an approximation of the original TRF. It is possible that the dynamic warping of more precise models of the original TRF could produce even more accurate EEG predictions. Second, in order to prevent overfitting, we chose to dynamically warp the TRF on every channel using a single set of transformation parameters. This may be suboptimal. Future work, likely needing larger datasets, might be able to learn channel specific modulations that would provide a more accurate model of the data. Third, due to the current group-level training protocol, the modeling we have carried out here cannot capture between-subject differences. It is natural to suppose that people have idiosyncratic neural responses, especially in challenging listening conditions. Training person-specific models may be a profitable avenue for future research. Finally, the module we used for estimating the transformation parameters was itself a linear convolutional network. As such, it may not have been able to capture what might be interesting non-linear relationships between the data and the transformation parameters.

## Methods

### Data preprocessing

The datasets we used are publicly available (Broderick et al., 2018). All EEG data were acquired at a rate of 512 Hz using an ActiveTwo system (BioSemi). 128 EEG channels plus two mastoid channels were used. The EEG data were re-referenced to the average of the mastoid channels, band-pass filtered between 0.5 and 8 Hz and downsampled to 64 Hz in MNE-Python (Gramfort et al., 2013). The noisy channels were detected by: (1) finding and excluding channels with EEG whose standard deviation (std) was 2.5 times larger than the averaged std of all channels, (2) finding and excluding channels (from all channels) with EEG whose std was 0.4 times smaller than the average std of the remaining channels, (3) combining channels found in (1-2) to get the noisy channels. We then interpolated those noisy channels using the spherical spline interpolation provided by MNE-Python. The obtained data was finally z-scored within each trial.

### Experimental procedure

In the single talker condition, each participant was instructed to listen an audiobook version of the “The Old Man and the Sea” read by a single male American speaker. In total, 19 participants took part in the experiment, with each listening to 20 trials lasting about 180 seconds each.

In the cocktail party condition, each participant was instructed to attend to one talker when the audiobooks “Journey to the Center of the Earth” and “20000 Miles Under the Sea” were simultaneously presented, one to each ear. The two audiobooks were read by two different male speakers. A total of 33 participants took part in the experiment, which consisted of 30 trials, each lasting about 60 seconds.

For both experiments, the segments of the audiobooks were presented in sequence, with each trial continuing from where the previous trial concluded, i.e., the storyline was preserved.

### Speech stimulus representation

As we were interested in assessing how EEG reflects the processing of the semantic processing of words, we needed to define a representation of the speech to relate to the EEG responses. Following other recent studies (Heilbron et al., 2022; Synigal et al., 2023), we chose to do this by computing a measure of lexical surprisal for each word.

### Lexical surprisal

Lexical surprisal quantifies how ‘surprising’ or how unexpected a word is. It is defined as the negative logarithm of a word’s probability of occurrence (here we used the natural logarithm):

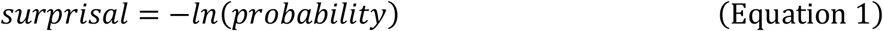

To estimate unexpectedness of a word given prior contextual information, we derived the lexical surprisal using the large language model Generative Pre-trained Transformers 2 (GPT-2).

As with other types of so-called transformer based models (Vaswani et al., 2017), GPT-2 is trained to predict the next word using previous context information (Radford et al., 2019). We restricted our use of GPT-2 to such a causal implementation in the absence of any strong evidence – as yet – that humans use predictions about future words to predict the current word. In practice, GPT-2 predicts the occurrence of a token instead of a word. A token is a common piece shared by words and is the minimal unit that a transformer model uses to model natural language. The surprisal of a token is calculated using its probability as estimated by GPT-2 (which is the conditional probability based on its prior tokens):

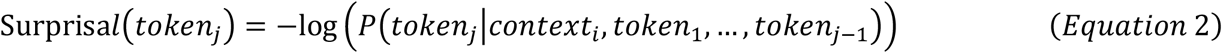

Here, we chose to incorporate the maximum number of tokens that GPT-2 can use, which is 1024. Thus, based on the chain rule in probability theory, lexical surprisal for each word is the sum of the surprisal estimates for all tokens making up that word:

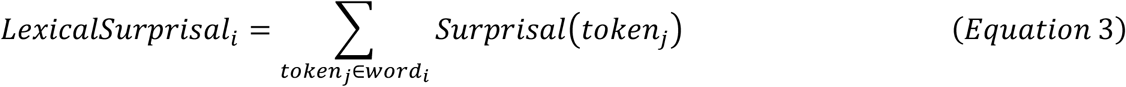

where *context*_*i*_ in Equation 2 indicates the context of *word*_*i*_, and {*token*_*j*_ ∈ *word*_*i*_| *j* = 1 … *N*}.

For each trial, the surprisal of the first token is set to zero. This ignores the fact that each audiobook segment continues from where the previous segmented ended. But, given the intertrial delays (with participants sometimes answering comprehension questions and/or taking breaks), we simply assumed zero context at the beginning of each trial. Punctuation marks were included in the input of GPT-2.

### Phonetic uniqueness point

The phonetic uniqueness point estimates the time point (relative to word onset) that the word is uniquely disambiguated from the lexical cohort of phonetic neighbors that begin with same phoneme sequence. To compute this time point, we first define what is known as the phonemic cohort entropy of each word.

The Phonemic Cohort Entropy of a word at the *i*th phoneme was derived using the following equation:

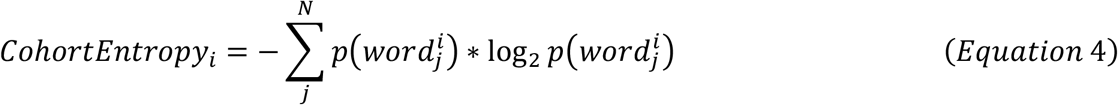

where N indicates the size of the cohort, and 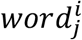 indicates the word cohort sharing the same phoneme sequence as the target word until the *i*th phoneme (Brodbeck et al., 2018). This measure decreases as more phonemes are heard (i.e., as fewer and fewer potential words remain available that are compatible with the phoneme sequence).

In an attempt to separate out within-word phonetic distinguishability from context-constrained word predictability, we chose here to use absolute word frequency to quantify word probability instead of the context-based predictability that derives from GPT-2. Specifically, the probability of the *j*th word in its cohort at the *i*th phoneme was derived as:

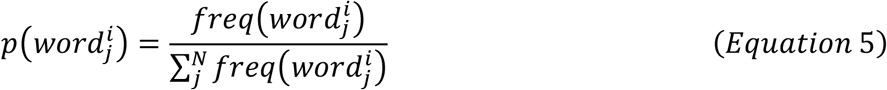

The word frequency information is obtained from the SUBTLEXus database (Brysbaert & New, 2009).

Having computed phonemic cohort entropy for each word (time aligned to the end of the corresponding phonemes in that word), we identified the uniqueness point as that point where the phonemic cohort entropy stopped decreasing.

### Word onset impulse

The timestamps of onsets for words in the audio book were obtained by applying a forced-alignment software Prosodylab-Aligner (Gorman et al., 2011) on the audio and the corresponding timestamps. A unit impulse was placed in a zero vector at the location corresponding to the timestamp of word onsets in terms of a sampling rate of 64. These impulses were included in the analysis to account for variance in the EEG that might arise at word onset unrelated to lexical surprisal.

### Lexical surprisal impulses

Sharing the same procedure as building the word onset impulses. However, instead of a unit impulse, the amplitude of the impulses here were the corresponding lexical surprisal values.

### Speech envelope

The broadband amplitude envelope was calculated using the Hilbert transform. As with the word onset impulses, this representation was included as a predictor in the TRF to absorb variance in the EEG that might arise from acoustic variations in the speech unrelated to lexical surprisal.

### The Dynamically Warped TRF

#### Overview

The core idea behind the proposed approach is to dynamically warp the static TRF with context-specific linear transformations (Figure 1B, 1C). Since it is difficult to manually design the formula for such a transformation, we estimated transformation parameters in a data-driven way using gradient descent (Curry, 1944). Thus, to update the weights that are used for generating transformation parameters, the process of transforming the static TRF with transformation parameters requires that that TRF be a differentiable function. Therefore, we here chose to convert the static TRF computed from the EEG data into a differentiable continuous function h (Figure 1A). Once represented in this way, this functional representation of the static TRF can be transformed into the dynamic TRF as follows:

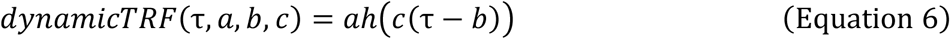

where τ are time lags referenced to word onset (as in a standard static TRF), *a* represents amplitude scaling, *b* represents time shift, and *c* represents time-scaling.

#### The static TRF

The dynamic-TRF begins with the computation of a static TRF before parameterizing the functional representation of this static template (Fig. 1A). A static TRF is a response function which describes the human brain as a memoryless linear time-invariant system in processing natural speech. The weights of it can be solved using a standard ridge regression mapping from time-lagged stimulus features (to account for time lags within the response function) to individual electrode responses (Crosse et al., 2016). To use the static TRF to predict unseen EEG, one can simply convolve the TRF with the time-series of the speech feature of interest (e.g., lexical surprisal at word onsets). The formula of the convolution operation is:

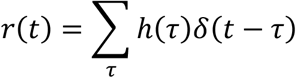

#### Functional Representation of Temporal Response Function

We built a functional representation (H) for the static TRF using functional data analysis, which decomposes the target TRF template vector into a linear combination of a group of continuous basis functions (Fig. 1A right). The discrete TRF template was estimated based on ridge regression using the mTRFpy library (Bialas et al., 2023). The continuous fit was performed using functional data analysis with Fourier basis (21 bases) using the scikit-fda library (Ramos-Carreño et al., 2019). The Pearson’s correlation between the functionally represented TRF and the original static TRF averaged across channels was ∼0.99.

#### The Parameterization of the Dynamic TRF Transformation

The parameters that were used to transform the functional representation of the TRF were estimated using an data-driven module built using PyTorch (Paszke et al., 2019). By default, the dynamic TRF has a causal convolutional layer having separate output nodes – one for each transformation parameter (Fig. 6).

**Fig. 6.**
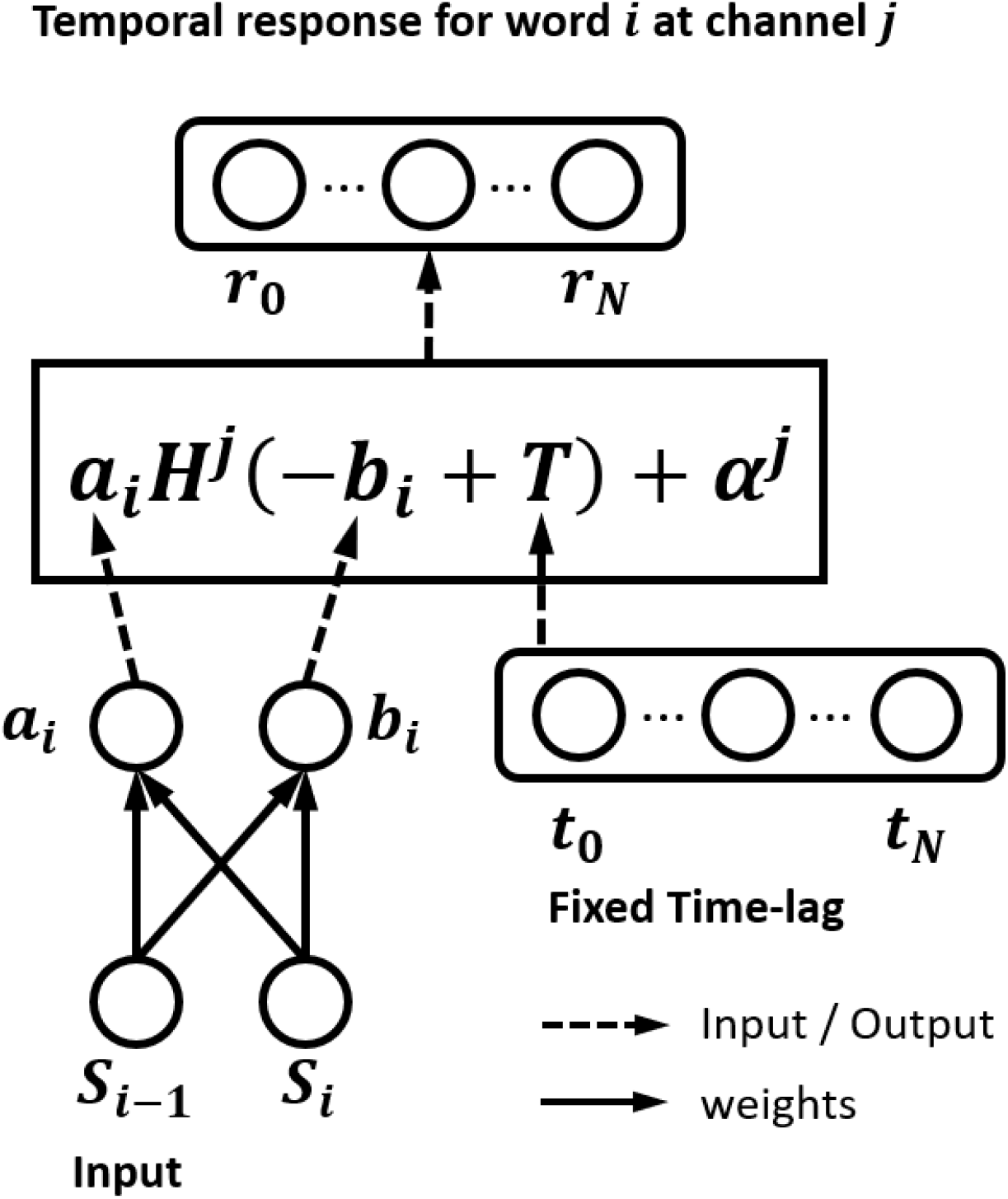
Details of the dynamically warped surprisal TRF model supporting time shifting and amplitude scaling with a window length of two. The causal convolutional layer (left bottom) takes the lexical surprisal (or phonetic uniqueness point) of the current and previous words as input to generate the parameters for time shifting (*b*_*i*_) and amplitude scaling (*a*_*i*_). The time-shift parameter was broadcast pointwise summed with a time lag embedding (*t*_0_ … *t*_*N*_). Given a TRF window between 0 ms to 700 ms, and a sampling rate of 64 Hz, *t*_0_ will be zero and *t*_*N*_ will be 45. The summed *b*_*i*_ and time lag embedding is fed into the function TRF H whose output is broadcast pointwise multiplied with *a*_*i*_ and broadcast pointwise summed with the interception of channel *j* (α^*j*^). The final dynamically warped temporal response (*r*_0_ … *r*_*N*_) has the same length as the time lag embedding.

When assessing whether or not modulating time shift and amplitude scaling could be accomplished with just a single variable (Fig 4), we built another dynamic TRF using a slightly different architecture. Specifically, the initial convolutional layer had a single output node, which provided the input to a new linear layer that had two separate nodes one for each transformation parameter. In this way, the module learned both an amplitude scaling parameter and a time-shifting parameter (in their appropriate units), but constrained them to be dependent on surprisal in the same way.

The weights of the causal convolutional layer were optimized on the same EEG training dataset from which the static TRF template was derived. For interpretability, the network was configured to have only six trainable weights (corresponding to the three TRF transformation parameters for the current and previous words). Weights for combining the Fourier basis set to get the 128 channels of the dynamically warped “N400” TRF were fixed during network optimization. Network loss was computed as the Mean Square error across all electrodes, computed between the entire time-series of predicted EEG data and the corresponding real EEG data. Training was achieved using the AdamW algorithm (Loshchilov & Hutter, 2018) in batches of size one run (e.g., 3min of an individual participant’s EEG data) repeated across 200 epochs (training stops early when the prediction accuracy stops increasing for 10 epochs). The learning rate was cycled between two boundaries (1e-3 to 1e-2) with the higher boundary reduced after each four epochs (a cycle) to reduce the difference between two boundaries by two (see Training Details section) which is referred as the “triangular2” cyclical learning rate policy (L. N. Smith, 2017). The weight decay (L2 regularization) was set to 1e-3. Error gradients were estimated automatically via torch.autograd.

To prevent overfitting, we used the same transformations on all time lags and channels. Thus, given the estimated transformation parameters (amplitude scaling *a*_i_, time shift b_*i*_, and time scaling c_i_, the TRF for the ith word at the jth channel is given by the following function (parameters colored in orange were fitted during gradient descent optimization):

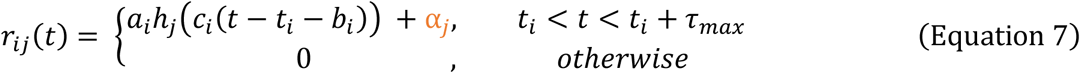

where α_*j*_ is the bias term of the jth channel of the dynamic TRF function, *t*_*i*_ indicates the onset time of the *i*th word, and:

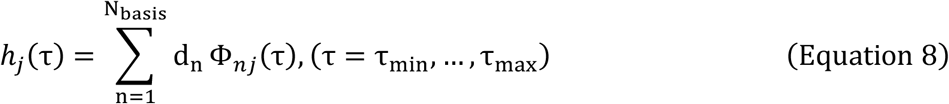

given a window length of *N* + 1, for the dynamic TRF without shared weights:

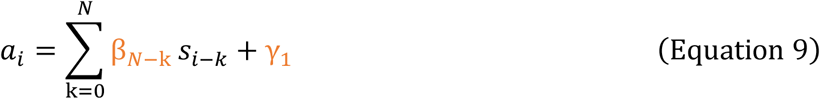

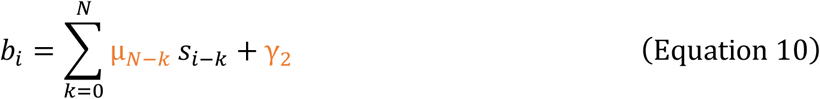

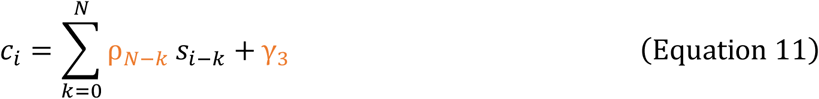

for the dynamic TRF with shared weights:

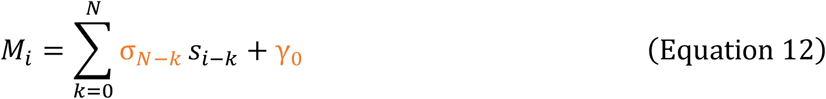

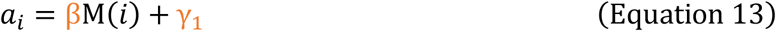

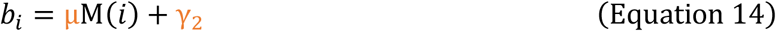

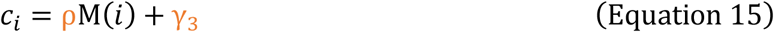

where *d* and Φ indicate the coefficient and function of the basis, respectively; σ, β, μ and ρ indicate the weights of the causal convolutional layers, with K indicating the length of convolution window and γ indicating the bias terms. We set the τ_*min*_ as 0 ms and τ_*max*_ as 700 ms. In the real implementation, we constrained the interval of time shift as -200 to 200ms, thus the static TRF that the dynamic TRF was built on was fitted with a minimum time lag of -200ms and a maximum time lag of 900ms. Each type of transformation parameter also had a default value. For word *i*, its default amplitude scaling parameter is the same as the static TRF, which is the predictor value of that word. The default time shifting is set to 0, and the default time-scaling is set to 1. The corresponding default value is used when the type of parameter is not enabled. Instead of using the convolution formula as shown in the above section of static TRF, here we represent the dynamically warped TRF function as the summation of temporal responses of words:

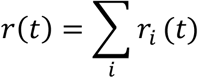

where *r*_*i*_(*t*) indicates the temporal response of the *i*th word.

### Accuracy Metric and Loss Function

The loss function we used to train the model was the mean squared error between the predicted EEG and true EEG. We measured the prediction accuracy by calculating the Pearson’s correlation between the predicted EEG and true EEG on each channel.

#### Cross validation

We used a standard 10-fold cross-validation procedure to test the dynamic TRF. Each dataset was equally split into 10 partitions, with each partition containing data from all the participants for 1/10 of the stimuli. Within each iteration of cross-validation, 8 partitions were used for training, 1 partition was used for validation, and 1 partition was used for testing the model. Training folds were used both to estimate the static TRF template and to train the dynamic TRF model. For the static TRF model, which is fitted using ridge regression, the validation set was used to choose the regularization hyperparameter (Crosse et al., 2016, 2021). For the dynamic TRF model, which is fitted using gradient decent, the validation set was used to both choose the regularization hyperparameter for the static TRF as well as decide when to stop training the dynamic parameters. Training was stopped when the prediction accuracy failed to increase for 10 epochs (an epoch indicates a single pass through the entire dataset). The partitions used for training, validation, and test were rotated across validation iterations. In detail, we first divided each story within each dataset (single talker, attended speaker, and unattended speaker) into 10 partitions. In the first round of cross-validation, we chose the EEG recordings corresponding to the first story partition as the testing set, the partition immediately subsequent as the validation set, and the remaining partitions as the training set. We did the same for the remaining 9 rounds of cross-validation, and if the last partition was used for testing, the first partition was chosen for validation.

#### Statistical Analysis

Comparisons between the EEG prediction accuracies of two models to be compared (e.g. the static TRF and dynamically warped TRF) were calculated using the Wilcoxon signed-rank test across subjects provide by the SciPy library (Virtanen et al., 2020). Since we selected the ‘exact’ method to calculate the signed-rank test statistics, which is recommended by SciPys when the number of samples is below 50, only W statistics were provided. Because the proposed model was hypothesized to improve the prediction accuracy, all the tests are right-tailed unless explicitly mentioned. When comparing the prediction accuracy for data on individual electrodes, the p-value obtained for each electrode were corrected using Benjamini-Hochberg correction-based false discovery rate provide by the Statsmodels library (Seabold & Perktold, 2010), with the alpha level set to 0.05.

## Supplementary Material

### Influence of Convolutional Window Length

In the analyses presented in the main body of the manuscript, we used a convolutional window of two when fitting all dynamic TRFs. This means that the causal convolution layer that learns the TRF amplitude, time-shifting and time-scaling parameters only uses the lexical surprisal values for the current word and the previous word. To validate that this was a reasonable choice, we tested if using other window lengths would improve the ability of the dynamic TRF to predict left out EEG data. We did this based on the Time&Amp dynamic TRF as this was generally the best performing model across our various datasets.

The extended Figure 1 shows the median prediction accuracy (averaged across all scalp channels) for the static TRF (dashed lines), and for the Time&Amp dynamic TRF (dots, solid lines). The error bars show the central 68% of the sampling distribution (equivalent to one standard error for a Gaussian). Errorbars were calculated using within-subject standard errors (Loftus & Masson, 1994), which removes variation across subjects that is shared between all conditions. Specifically, before calculating the central interval, the mean for each subject across all conditions was removed. A correction factor was added to the interval to make the estimation unbiased. The correction factor was calculated as 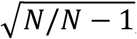, where *N* indicates the number of conditions. In the single talker condition, improvements started to be significant for a time window of two, however, the improvement in prediction accuracy for a window length of 3 over 2 was not significant (p = 0.0668, Wilcoxon Signed-rank Test). In the attended speech condition, only the time window of 2 had significant improvement over the fixed TRF. In the unattended speech condition, there was no significant improvement for any window length. These results suggest that the lexical surprisal for the current and previous words are the most influential in modulating the amplitude and time latency of the N400 TRF.

**Supplementary Fig. 1.**
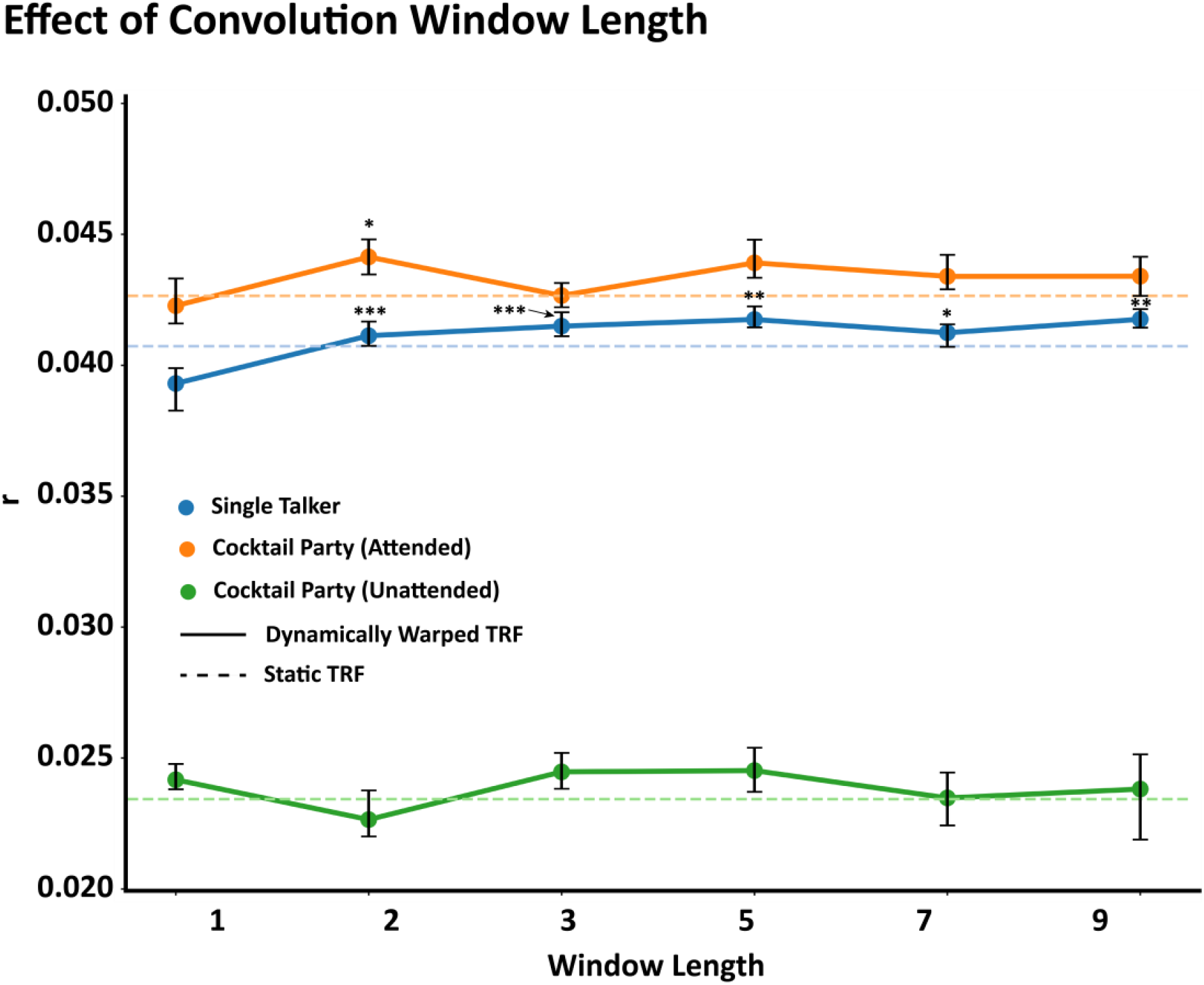
The effect of the window length used to predict amplitude scaling and time-shift parameters. This figure plots the prediction accuracy (the median across subjects) for the dynamic vs. the fixed TRF models as a function of the window length used to predict the amplitude scaling and time-shift parameters of the variable TRF. Results are shown separately for the single talker, attended and unattended conditions. The dashed line indicates the performance for the static TRF. For single talker and attended speech, the model achieved the highest prediction accuracy when its window size was around 2-3 (the improvement from 3 to 2 for single talker was not significant, p = 0.0668). For unattended speech, the prediction accuracy was always not significant compared with the accuracy of the static TRF.

**Supplementary Fig. 2.**
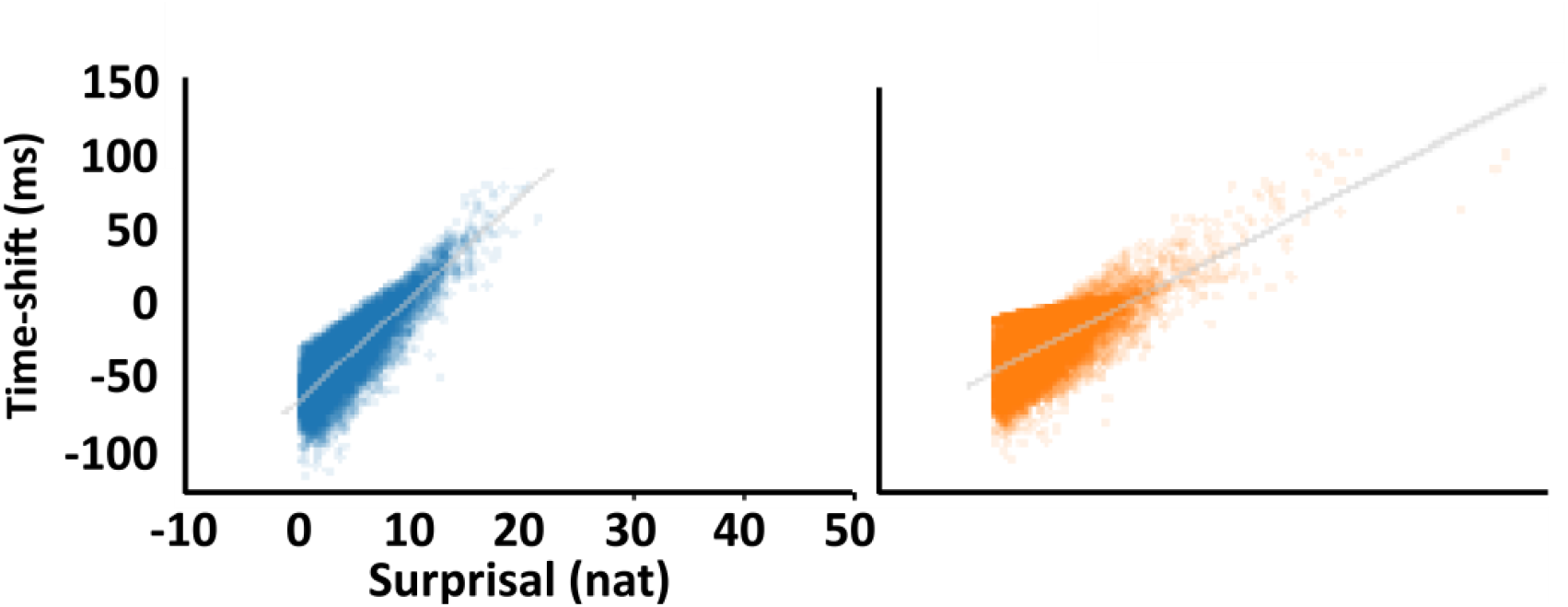
A, The scatterplot for surprisal and the time-shift. Each point indicates a pair of these two variables of a word. B, The scatterplot for uniqueness point and the time-shift. Each point indicates a pair of these two variables of a word.

